# Biologically Plausible Neural Networks for Simulating Brain Dynamics and Inferring Connectivity

**DOI:** 10.1101/2024.12.20.629853

**Authors:** Aarush Gupta

## Abstract

We present Cerebrum, a novel framework that bridges biologically plausible neural networks with rigorous mathematical modeling. By grounding neural network simulations in a range of empirically-founded neuronal models (Hodgkin–Huxley, Izhikevich, and Adaptive Exponential Integrate-and-Fire), Cerebrum captures dynamics from intricate ion channel kinetics to general, large-scale network behavior. Our methodology integrates these mathematical models within a unified simulation pipeline, featuring mechanisms for short-term synaptic plasticity, precise state integration via the Runge-Kutta and Euler methods, and refractory spike regulation. We use a Graph Attention Network (GAT), to infer synaptic connectivity directly from dynamic activity patterns. This approach not only ensures biological realism but also facilitates robust connectivity reconstruction across diverse network topologies, including Erdős–Rényi, Small-World, and Scale-Free architectures. Results demonstrate that our framework reliably reproduces key neuronal firing patterns and synchronization phenomena, while its modular design paves the way for scalable investigations into the interplay between network structure and neural dynamics. Cerebrum thus offers a powerful tool for advancing both theoretical and applied aspects of computational neuroscience.

## 1 Introduction

Accurate simulation and inference of brain network dynamics remain major challenges in computational neuroscience. Traditional graph-theoretic analyses have elucidated many topological features of neural systems [16], yet they often neglect the intrinsic temporal dynamics of neuronal activity. In this work, Cerebrum introduces a paradigm shift by integrating biologically detailed neuronal models with state-of-the-art connectivity inference techniques. The framework not only advances simulation fidelity through a multi-model approach—encompassing the Hodgkin–Huxley, Izhikevich, and Adaptive Exponential Integrate-and-Fire models—but also enhances interpretability via a Graph Attention Network that reconstructs synaptic connectivity directly from dynamic activity patterns. This innovative integration addresses longstanding challenges by offering a unified, scalable platform that bridges detailed neurodynamics with robust network inference, ultimately opening new avenues for understanding the interplay between neuronal behavior and network topology. The novelty of Cerebrum lies in its modular design, its capacity to simulate diverse network architectures, and its ability to incorporate disease-specific modifications, all of which collectively push the boundaries of current computational neuroscience methodologies.

The central problem we address is twofold: (i) how to accurately simulate the dynamic behavior of neurons using both biophysically detailed and computationally efficient models and (ii) how to infer the underlying synaptic connectivity from the resulting activity patterns in a robust, interpretable, and adaptable manner. By integrating multiple neuronal models with a GAT, Cerebrum offers a unified, modular platform for simulating, analyzing, and optimizing brain network dynamics—an approach that resonates with contemporary findings in both dynamical neuroscience and network analysis [14].

## 2 Methodology

### 2.1 Topologies and Data Sources

To generate ground-truth connectivity matrices, Cerebrum supports both real connectome data (e.g., *C. elegans* synaptic matrices) and synthetic networks. In the absence of empirical data, synthetic connectivity is generated based on canonical network topologies such as Erdős-Rényi [5], Small-World [20], ScaleFree [13], and Modular Small-World architectures. When synthetic, connectivity is determined by a connection probability parameter *p*, ensuring control over the average degree 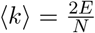 for fair cross-topology comparisons.

### 2.2 Neuronal Models

Cerebrum implements several mathematical models to simulate neuronal dynamics, thereby catering to varying requirements of biological accuracy and computational efficiency. In our framework, the detailed Hodgkin-Huxley (HH) model is employed to capture ion channel dynamics, as governed by the differential equation

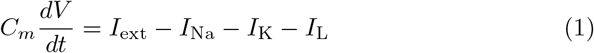

with ionic currents defined as

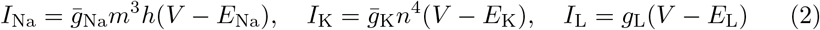

Gating variables follow first-order kinetics:

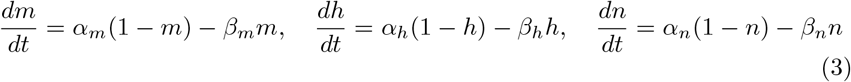

For computational efficiency, the Izhikevich model [11] and the Adaptive Exponential Integrate-and-Fire (AdExp) model [1] are also implemented. This strategy optimally balances biophysical fidelity with computational cost, reflecting the empirical trade-offs observed in large-scale neural simulations [7].

### 2.3 Simulation

The simulation pipeline in Cerebrum is organized into discrete time steps and comprises several tightly coupled modules. At each time step, the following sequence is executed:

### 1. Short-Term Plasticity (STP) Update

For a synapse from neuron *j* to neuron *i*, we maintain two dynamic variables *uij*(*t*) (facilitation) and *xij*(*t*) (depression). Their dynamics, inspired by the Tsodyks–Markram model [18], are updated in discrete time with step Δ*t*. When a presynaptic spike is detected at time *t* (i.e., *Sj*(*t*) = 1), the updates are

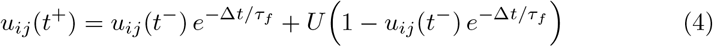

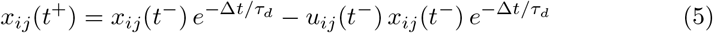

where *τf* and *τd* are the time constants for facilitation and depression, respectively, *U* is a model parameter, and *t*− and *t*+ denote the instants immediately before and after the spike. In the absence of a spike (*Sj*(*t*) = 0), the variables relax as

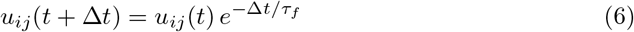

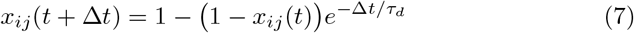

The effective synaptic weight is computed as

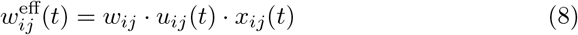

with *wij* being the baseline synaptic weight.

### 2. Synaptic Current Computation

A GAT module computes the synaptic current for neuron *i* at time *t* by aggregating contributions from its presynaptic neurons:

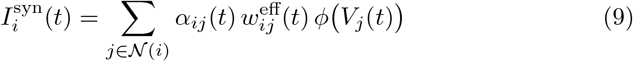

where 𝒩 (*i*) is the set of presynaptic neurons, *Vj*(*t*) is the membrane potential of neuron *j*, and *ϕ*(·) is an optional nonlinearity. The attention coefficients *αij*(*t*) are defined as

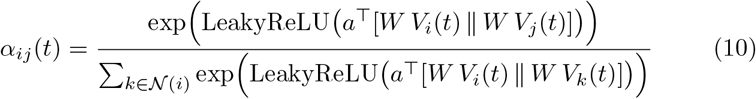

with *W* as a learnable weight matrix, *a* as a learnable attention vector, and ∥ denoting concatenation. For instance, in a GAT, the attention coefficients are computed as

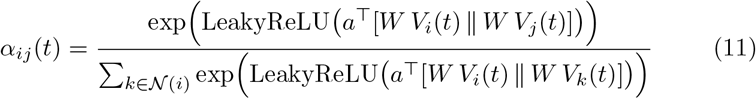

### 3. Synaptic Delay Buffer

Synaptic transmission delays are modeled via a delay buffer. The synaptic current arriving at neuron *i* at time *t* is given by

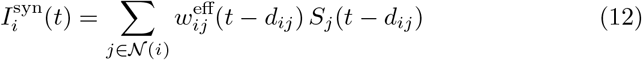

where *dij* is the delay for the connection from neuron *j* to *i*, and *Sj*(*t*−*dij*) is the spike indicator at the delayed time.

### 4. State Integration

Let **y**(*t*) denote the state of a neuron (e.g., membrane potential and gating variables). Its evolution is governed by the first-order ordinary differential equation

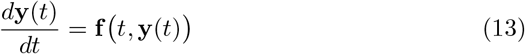

We use the fourth-order Runge-Kutta integration method [3] for the HH model and the Euler integration method [8] for the Izhikevich and AdExp models.

### 5. Spike Detection and Refractoriness

Spikes are detected by threshold crossing of the membrane potential:

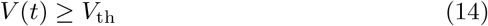

where *V*th is the threshold. The spike indicator is defined as

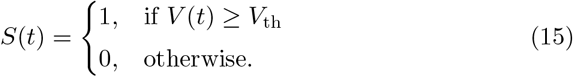

A refractory period *T*ref is enforced (e.g., by clamping *V* (*t*) or inhibiting further updates) during the interval *t* ∈ [*t*spike, *t*spike + *T*ref].

### 2.4 Connectivity Inference

To recover the underlying synaptic connectivity from the simulated activity, Cerebrum employs an GAT. The model processes node-level features (e.g., mean membrane potentials, firing rates, and voltage variances) and reconstructs the adjacency matrix Â. For each potential connection (*i, j*), the inferred weight is computed as

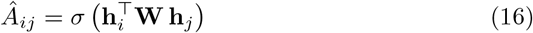

where **h***i* and **h***j* are learned node embeddings, **W** is a weight matrix, and *σ* is the sigmoid activation. The network is trained to minimize the mean squared error (MSE) between the predicted and ground-truth matrices, using gradient clipping and dropout-based uncertainty estimation.

### 2.5 Modifications and Optimization

Cerebrum further enables the simulation of pathological conditions by modulating neuronal parameters. For example, Parkinson’s-like dynamics are modeled by increasing inhibitory tone and reducing external excitatory currents, while Epilepsy-like conditions are simulated by enhancing excitability and promoting synchronized bursting [9, 12]. These modifications are applied via a specialized Neuron Editor module that scales parameters based on a disease configuration. Parameter optimization is performed by exploring a predefined parameter space. Short simulation runs yield performance metrics (e.g., firing rate, synchronization, and variability), and the configuration with the optimal score is selected. This systematic approach is consistent with recent studies emphasizing robust parameter estimation in brain network models.

## 3 Results

Our evaluation of Cerebrum spans multiple network topologies and neuronal models. In synthetic networks comprising 200 neurons with an average connection probability of 0.05, the Hodgkin-Huxley (HH) model [10] produced dynamics that closely resembled biological observations when paired with the attention-based connectivity inference module [19]. Figures 1 and 2 illustrate the firing rate distributions and membrane potential traces for Erdős-Rényi, Small-World [20], and Scale-Free networks. Notably, Scale-Free networks exhibited lower firing rates (0.006 spikes/unit) and reduced synchronization (6.19) compared to Erdős-Rényi (0.053, 11.10) and Small-World (0.056, 13.96) networks, indicating that SF networks yield clearer, less-confounded neural signals, thereby facilitating more accurate connectivity inference [2].

**Figure 1:**
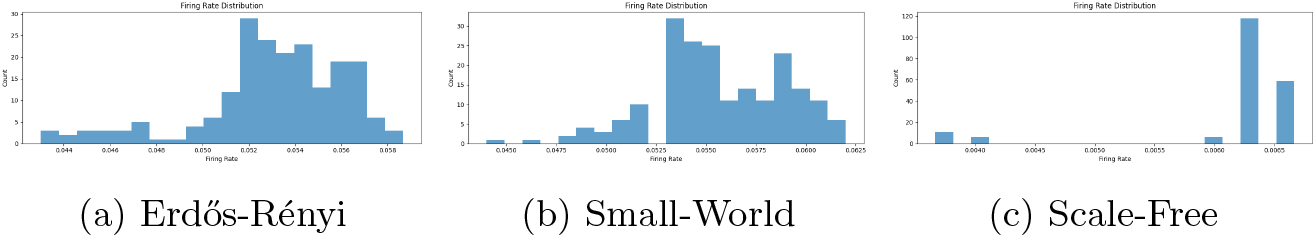
Firing rate distributions across network topologies. Scale-Free networks exhibit lower and more selective firing rates compared to Erdős-Rényi and Small-World architectures.

**Figure 2:**
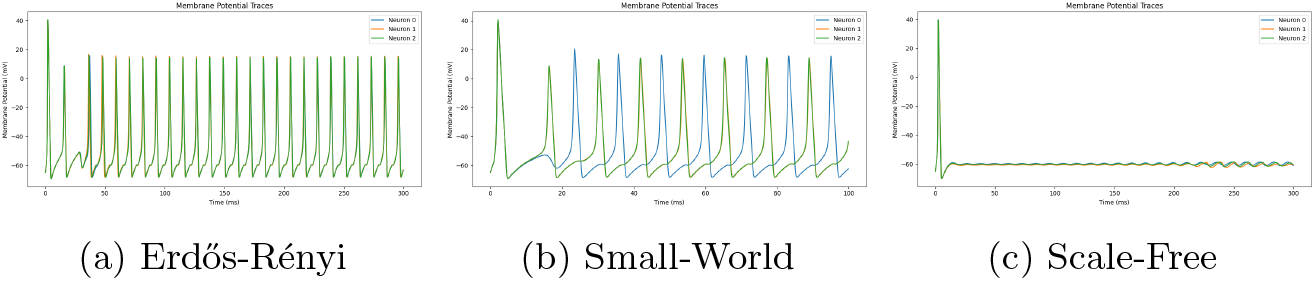
Membrane potential traces across network topologies. The Scale-Free network demonstrates sparser and less synchronized activity compared to other topologies.

The structural differences among network topologies significantly influenced neuronal firing activity and synchronization patterns. We observed that the Scale-Free network exhibited markedly lower spiking rates and reduced synchronization compared to the ER and SW networks, as detailed in Table 1, aligning with observed biological phenomena [4, 15].

**Table 1:** Comparison of spiking rate and synchronization for different network topologies.

| Network Topology | Spiking Rate | Synchronization |
| --- | --- | --- |
| Erdős-Rényi | 0.053 | 11.100 |
| Small-World | 0.056 | 13.960 |
| Scale-Free | 0.006 | 6.187 |

Functional connectivity matrices further revealed that the heterogeneous correlation structure in SF networks provided richer discriminative features, thereby improving the accuracy of connectivity inference [2]. Disease-specific experiments demonstrated the framework’s versatility: while Parkinson’s-like modifications improved inference by sparsifying activity patterns [17], Epilepsy-like modifications led to overlapping activity that confounded the inference process [12].

Exploration of the parameter space revealed that Scale-Free networks are considerably more robust, with an average sensitivity of approximately 0.047 compared to 0.212 in Erdős-Rényi and 0.207 in Small-World networks. Dropout-based uncertainty estimation during inference resulted in nearly zero standard deviation across repeated predictions, indicating high confidence in the model’s connectivity outputs [6].

To further elucidate the performance of the connectivity inference module in Scale-Free networks, we present a side-by-side comparison of the predicted and ground-truth adjacency matrices (see Figure 3). The predicted adjacency (left) appears darker and exhibits a nearly grid-like pattern with pronounced horizontal and vertical lines, especially concentrated in the top-left region. This pattern likely reflects the network’s inherent clustering and the GAT’s focus on high-connectivity hubs. In contrast, the ground-truth adjacency (right) is primarily clustered in the top left, yet shows well-distributed, sporadic connections throughout the matrix. It is important to note that the color scales differ between the two heatmaps—the true adjacency ranges from 0 to 1 while the predicted adjacency spans from 0 to 2—which shows that the learned connectivity profile diverges in magnitude from the empirical ground truth. These insights, detailed further in Table 2, confirm that the connectivity inference module captures essential structural features of Scale-Free networks, even if some scaling adjustments are necessary to fully align the predicted weights with the true values.

**Table 2:** Comparison of parameter sensitivity and uncertainty for different network topologies.

| Network Topology | Avg. Sensitivity | Mean Uncertainty |
| --- | --- | --- |
| Erdős-Rényi | $\sim 0.212$ | 0.0000 |
| Small-World | $\sim 0.207$ | 0.0000 |
| Scale-Free | $\sim 0.047$ | 0.0000 |

**Figure 3:**
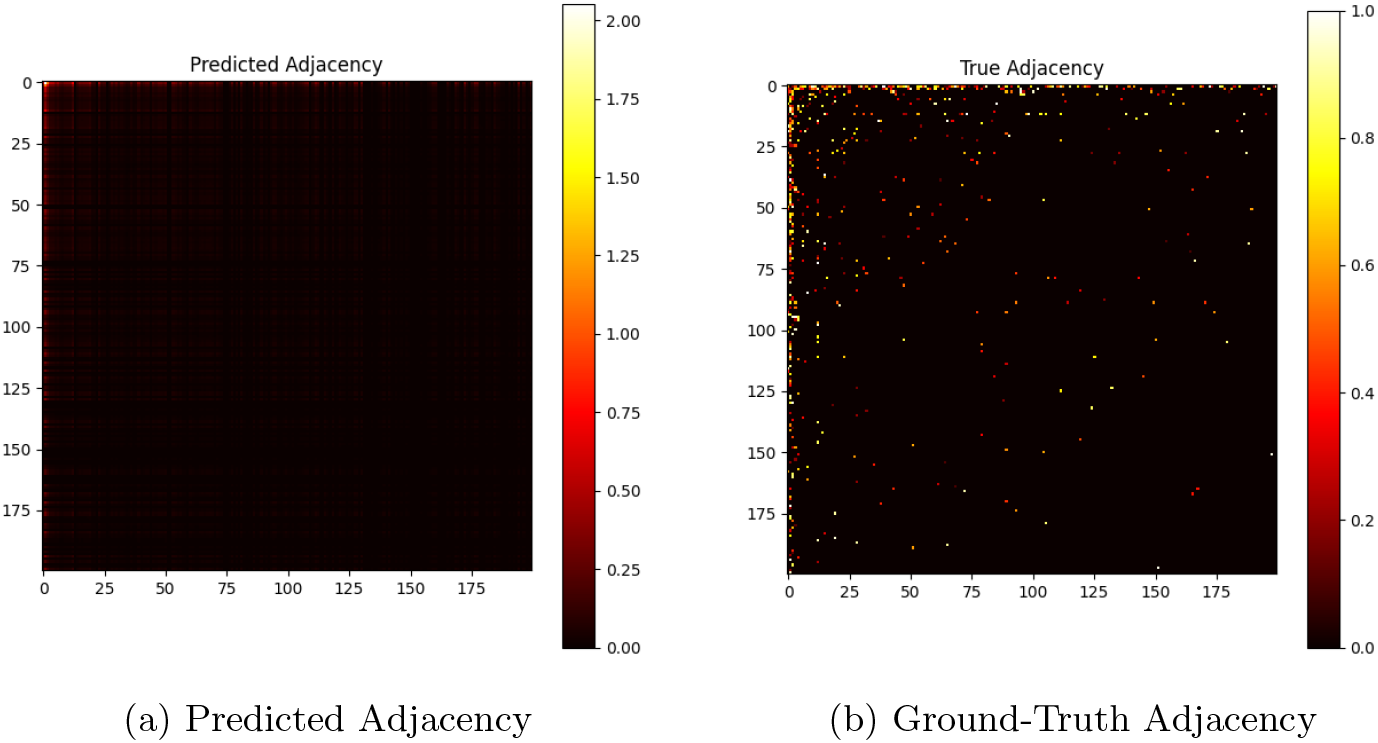
Adjacency matrix heatmaps for a Scale-Free network. The predicted adjacency (left) appears darker with grid-like horizontal and vertical lines, especially in the top-left region. This may result from compression effects or inherent scaling differences (predicted scale: 0–2 versus true scale: 0–1). The ground-truth adjacency (right) is primarily clustered in the top left with sparse connections elsewhere.

## 4 Conclusion

We have presented Cerebrum, a comprehensive and accessible framework for simulating biologically plausible neuronal dynamics and inferring synaptic connectivity. By integrating detailed and simplified neuronal models with a GAT for connectivity reconstruction, our system bridges the gap between biophysical realism and computational efficiency. The modular design—featuring dedicated modules for configuration, neuronal modeling, integration, synaptic plasticity, and connectivity inference—ensures that our codebase is both maintainable and extensible. Our simulation pipeline facilitates systematic exploration and optimization of brain network behavior. Our results indicate that networks with heterogeneous architectures, such as Scale-Free graphs, yield robust connectivity inference and dynamic patterns that closely mimic empirical observations. We anticipate that further scaling and refinement, will enable even more detailed studies of large-scale brain networks.

